# Multimodal Validation of the Existence of Transitional Cerebellar Progenitors in the Human Fetal Cerebellum

**DOI:** 10.1101/2025.06.01.657310

**Authors:** Zaili Luo, Mingyang Xia, Feng Zhang, Dazhuan Xin, Rohit Rao, Karrie M. Kiang, Kalen Berry, Yu Xiong, Hongqi Liu, Yifeng Lin, Ming Hu, Mei Xin, Jie Ma, Hao Li, Michael D. Taylor, Wenhao Zhou, Q. Richard Lu

## Abstract

The developing human cerebellum comprises a series of transient progenitor states that are essential for generating diverse neural subtypes, yet the identity and validation of intermediate cell populations bridging stem-like and lineage-committed neuronal precursors remain limited. In our previous single-cell transcriptomic study, we identified a distinct transitional cerebellar progenitor (TCP) population enriched in specific progenitor zones such as the rhombic lip during human fetal cerebellar development. To address the concerns raised in the *Matters Arising* regarding the existence of the TCP cells, we provide additional multimodal validations of this population. Rigorous reanalysis of our single-cell transcriptomic data, applying stringent quality control measures, validated the quality of TCP cells and their classification as a transcriptionally distinct population. Multiple orthogonal validations of TCP signature genes (*SOX11* and *HNRNPH1*) using RNAscope *in situ* hybridization, Xenium-based spatial transcriptomics, and immunohistochemistry on additional fetal cerebellar samples across different stages demonstrated the consistent presence of TCPs in the rhombic lip, transitioning from *PRTG*^+^ stem-like zones in the ventricular zone at early developmental stages to the subventricular zone overlapping with *EOMES*^+^ unipolar brush cell precursors at later stages. TCP-like populations were also independently identified in two fetal cerebellar single-nucleus transcriptomic atlases, and their gene signature was enriched in a cell population associated with aggressive medulloblastomas. Collectively, these multimodal validations confirm the existence of a transitional progenitor population in the human fetal cerebellum, with implications for cerebellar lineage progression and medulloblastoma origin.

## Introduction

The development of the human cerebellum is a highly coordinated and complex process that involves not only the sequential and concurrent generation and specification of distinct progenitor populations, but also their spatial patterning, migration, and integration into evolving neural circuits^1,2^. Despite significant progress in understanding cerebellar development, the transitional states bridging early neuroepithelial progenitors and lineage-restricted neuronal precursors remain poorly defined. In contrast to post-mitotic neuronal cell types, which exhibit discrete and stable marker expression patterns, transitional progenitor intermediates are often characterized by graded, quantitative shifts in the relative expression of a combinatorial marker signature, reflecting their dynamic progression along a developmental continuum^3-5^.

In our previous single-cell transcriptomic profiling analysis of fresh human fetal cerebellar tissues with commonly used filtering criteria and quality control measures, we identified a previously uncharacterized transitional cerebellar progenitor (TCP) population that exhibited unique transcriptional features distinct from known neural stem cells or committed neuronal lineage cells^6^. Immunostaining of human fetal cerebellar sections indicated that these TCP cells are spatially present in the progenitor-rich zones including the ventricular and transitional zones of the cerebellar anlage, including the rhombic lip, a critical germinal niche in the early developing human fetal cerebellum.

In April 2023, *Nature* sent us a *Matters Arising* “*Lack of evidence for the transitional cerebellar progenitor*” by Smith and colleagues^7^. To address the concerns raised, we present further evidence to validate the existence of this cell population in the developing fetal cerebellum. *We have 1) reconfirmed the quality of previously identified TCP cells with more stringent criteria, 2) validated the presence of TCP cells through different orthogonal methods by fluorescent in situ RNAscope, Xenium-based spatial transcriptomics, and immunohistochemistry assays of additional tissue specimens, 3) identified TCP-like signature genes highly expressed population in two independent fetal cerebellar single-nucleus transcriptomic datasets*^8,9^. We thank the *Matters Arising* authors for highlighting technical challenges and potential pitfalls with work on human cerebellar development. However, this reanalysis of our original data and additional experimental results confirm our previous conclusion regarding the presence of a TCP cell population in specific regions of the human fetal cerebellum such as the rhombic lip.

Our complementary approaches consistently identified TCP cells in the specific progenitor zones such as the rhombic lip, marked by TCP-signature gene expression in the ventricular zone during early fetal development and a shift to the subventricular zone at later fetal stages. Taken together, our additional analyses and orthogonal validations reinforce our previous findings identifying the TCP population as a bona fide transitional progenitor state within specific neurogenic geminal zones, such as the rhombic lip, during human fetal cerebellar development.

### Re-analysis of single-cell transcriptomics validates the data quality for the TCP population

In our original study^6^, by performing computational analyses of whole-cell single-cell transcriptomics data from fresh fetal cerebellar tissues, we identified the TCP population, which is enriched in the post-conception week (PCW) 12 specimen. This analysis utilized the standard pipelines implemented in CellRanger and Seurat with commonly used filtering criteria and quality control measures as well as doublet removal^6^. The *Matters Arising* claimed that our quality control (QC) filtering criteria and the use of the standard Seurat scRNA-seq analysis are liable to “result in the retention of more cells at the cost of imposing technical artifacts, such as the presence of ambient RNA”. To address potential concerns regarding ambient RNA contamination or low data quality-driven cell identification, we conducted additional analyses using more stringent quality control measures. These analyses employed established pipelines CellRanger and Seurat, alongside specialized tools including DoubletFinder^10^ and SoupX^11^, which are designed to detect and remove doublets and ambient RNA contamination from droplet-based scRNA-seq data. These rigorous filters re-confirmed that the TCP cells consistently passed all major QC thresholds, supporting the robustness and validity of their identification.

To further assess cell quality, we carried out additional DropletQC^12^ analysis, as requested by a reviewer, to enhance the identification of empty droplets and damaged cells in single-cell RNA-seq data. We first performed a nuclear fraction analysis of the TCP cell population in the PCW12 sample using DropletQC. Kernel density estimation supported the use of a nuclear fraction cutoff of 0.1 as an appropriate threshold to distinguish low-quality cells from high-quality ones (**Fig. 1a**). The vast majority of TCP cells (95.2%) in the PCW 12 sample exceeded the nuclear fraction cutoff of 0.1 (**Fig. 1b**), confirming their quality and arguing against the claim that they represent low-quality cells or ambient RNA contamination. This analysis directly contradicts the statement in the *Matters Arising* that “*Application of DropletQC [5] to the isolated PCW12 profile determined that TCP cells had a low nuclear fraction of detected transcripts*”. Even under a relatively permissive QC threshold (gene count >200) applied in the *Matters Arising*, the majority of TCP cells were still not classified as empty droplets (MA-Fig. 1g). We therefore conclude that this TCP cell population is of high quality, meeting the rigorous filtering criteria and DropletQC quality controls, and should not be excluded from bioinformatics analysis.

**Figure 1.**
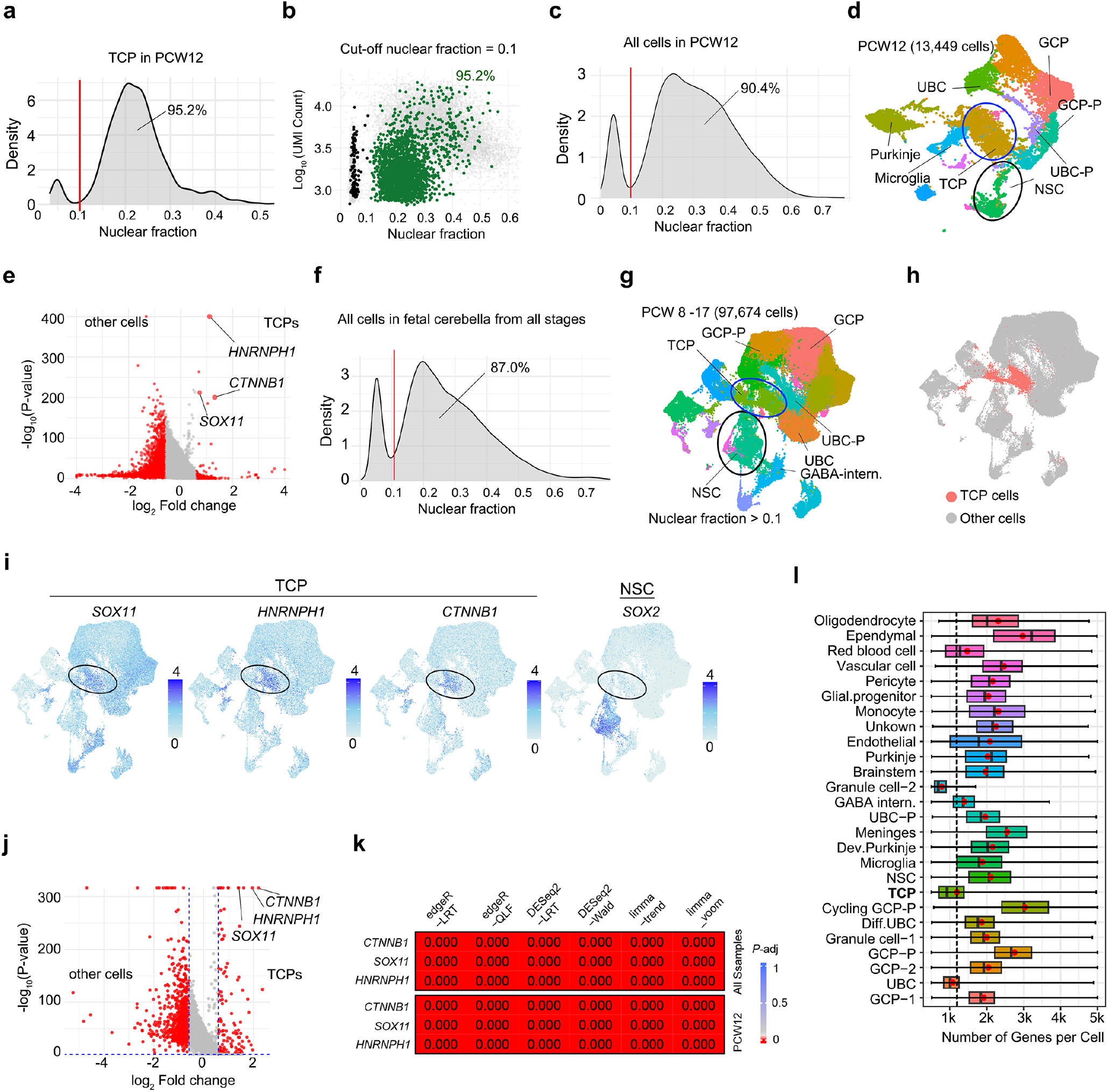
Additional quality controls including DropletQC re-confirm the presence of the TCP population in human fetal cerebella. **a**, Kernel density estimation for TCP cells in the PCW 12 sample by DropletQC. Red line, nuclear fraction cutoff at 0.1. **b**, Scatter plots of nuclear fraction and UMI counts per cell in the TCP cells from PCW 12 using indicated cutoff scores. **c**, Density plot of nuclear fraction for all cells in the PCW 12 sample. The red line, nuclear fraction at 0.1. **d**, UMAP plots of scRNA-seq data at PCW 12 using more stringent filtering criteria (>500 and <5000 genes and nuclear fraction >0.1). Circles, NSC and TCP populations. **e**, Volcano plots of differentially expressed genes (red, fold change >1.5) in TCP cells versus non-TCP cells at PCW 12 using a Wilcoxon rank sum test. **f**, Density plot of nuclear fraction for all cells in all samples of human fetal cerebella from PCW 8 to PCW 17. The red line, nuclear fraction 0.1. **g**-**h**, UMAP plots of human fetal cerebellar cells from PCW 8 to PCW 17 with more stringent filtering criteria and nuclear fraction cut-off > 0.1 with g) all cell types labeled and h) TCP distribution shown. Circles, NSC and TCP populations. **i**, Expression of TCP signature genes and neural stemness gene *SOX2* in UMAP plot. Circles, TCP populations. **j**, Volcano plot of differentially expressed genes (fold change >2) in TCP versus non-TCP cells with significance determined using a Wilcoxon rank sum test. **k**, Significance of differential expression of the TCP signature genes determined using the indicated statistical models to analyze pseudo-bulk data of the samples from all stages and at PCW 12. **l**. Boxplots summarizing the number of genes expressed per cell across cerebellar cell types (cells with > 500 or < 5000 expressed genes; nuclear fraction cut-off > 0.1). Boxes represent the first, median, and third quartiles; whiskers extend to 1.5 x the interquartile range. Red dots indicate the mean number of expressed genes per cell for each cell type. Dash line: mean number of expressed genes per cell in TCPs.

To more rigorously evaluate the cell quality of the PCW 12 sample, we reanalyzed these data using widely used filtering cutoff criteria^13-17^, which are supported by a machine learning framework^18^, namely, removing cells with < 500 or > 5000 expressed genes and > 20% mitochondrial gene content, together with DropletQC nuclear fraction cutoff of 0.1 determined through density estimation (**Fig. 1c**). Under these filtering criteria, unsupervised clustering identified TCP cells as a distinct population located adjacent to *SOX2*^+^ neural stem cells (NSCs) (**Fig. 1d**). Cells in the TCP cluster have significantly higher expression of TCP signature genes than do all other cells (**Fig. 1e**). Similarly, density estimation across all human fetal cerebellar cells from PCW 8 to PCW 17 supported a consistent nuclear fraction cutoff of 0.1 as appropriate (**Fig. 1f**). Applying these stringent QC filtering criteria across all samples, we consistently detected TCPs as a distinct cell population near NSCs through unsupervised clustering (**Fig. 1g,h**), with the significant elevation of TCP signature gene expression as determined using a Wilcoxon rank sum test in our original study (**Fig. 1i,j**). Notably, the differential expression of TCP signature genes was also demonstrated in the MA-Extended Data Fig. 2c of *Matters Arising* using the Wilcoxon rank sum test. Moreover, evaluating differential gene expression using additional six top-performing pseudo-bulk models of likelihood tests^19^, edgeR−LRT, DESeq2−LRT, edgeR−QLF, limma−trend, DESeq2−Wald, and limma−voom, we further confirmed significant differential expression of TCP signature genes (e.g., *SOX11, HNRNPH1, CTNNB1*) in TCP cells compared to non-TCP cell types at PCW 12 and at combined stages (**Fig. 1k**). These analyses refute the assertion in the *Matters Arising* that *‘zero significant marker genes’* exist in TCP cells. Together, the re-analyses of our original data confirm that TCP cells possess a distinct transcriptional identity characterized by a set of significantly enriched marker genes. Notably, under the stringent filtering criteria (requiring cells to express >500 and <5000 genes and a nuclear fraction >0.1), TCP cells exhibit a higher average gene expression (1,193 genes per cell) compared to well-defined post-mitotic unipolar brush cells (UBCs) (1,080 genes per cell) and granule cells (784 genes per cell), supporting the biological relevance of the transcriptional profile observed in the TCP cells (**Fig. 1l**).

**Figure 2.**
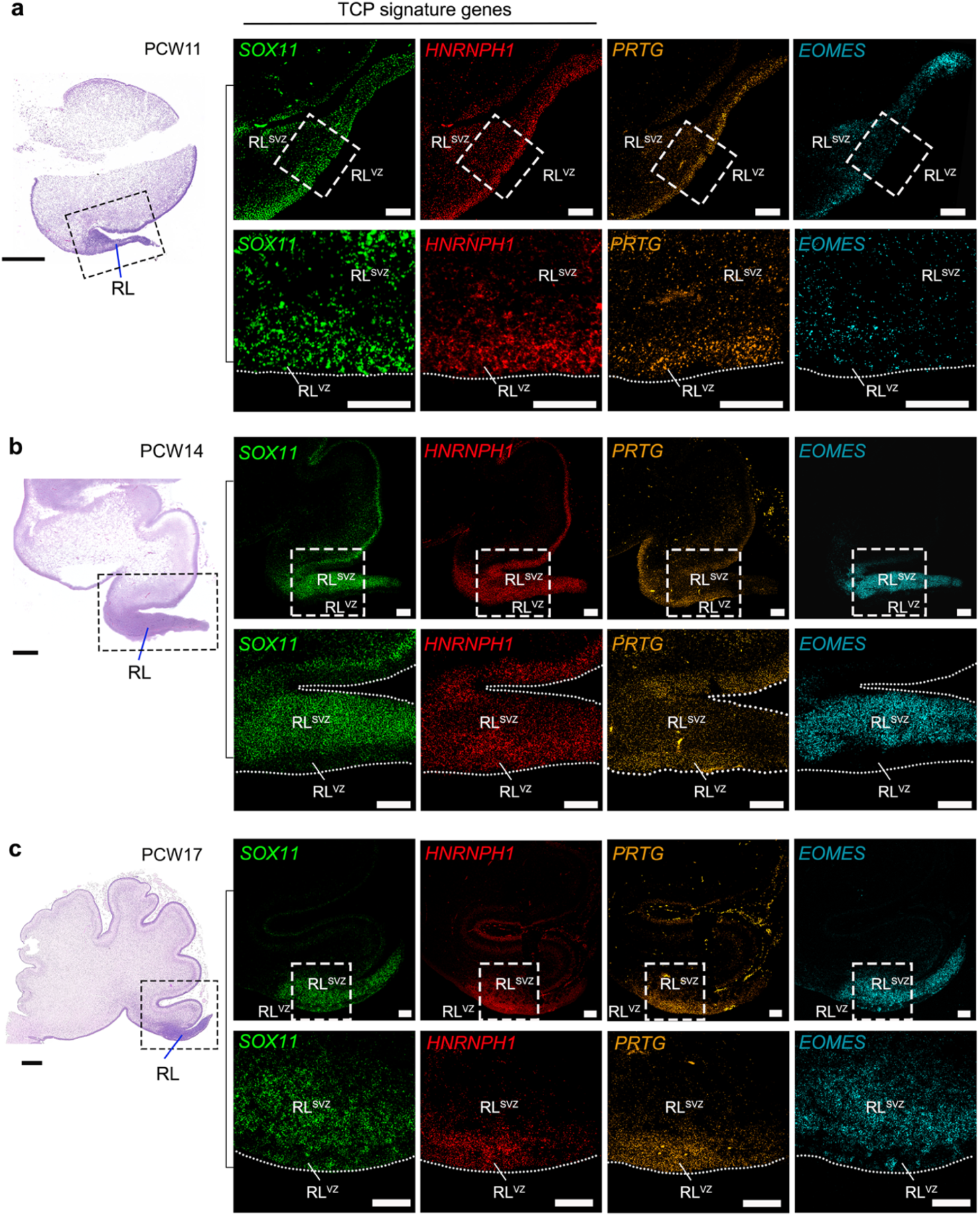
Orthogonal RNAscope fluorescent mRNA *in situ* hybridization validates the existence of the TCP population in the rhombic lip in the human fetal cerebellum. **a**-**c**) Left, hematoxylin/eosin staining of midsagittal sections in the human fetal cerebellum at post-conception weeks (PCW)11 (a), PCW14 (b), and PCW17 (c). Boxed areas: the rhombic lip (RL). Scale bars, 500 μm. Right, RNAscope fluorescent mRNA labeling of TCP signature genes, *SOX11, HNRNPH1*, and a stem-like cell marker *PRTG*, along with a UBC neuronal progenitor marker *EOMES* in the rhombic lip at PCW11, PCW14, and PCW17 as indicated. The boxed areas are shown in higher magnification in the corresponding lower panels. Scale bars, 100 μm. RNAScope probes: *SOX11* (Advanced Cell Diagnostics, Cat#443871-C2), *HNRNPH1* (Cat#1283271-C1), *PRTG* (Cat#1096421-C3), and *EOMES* (Cat# 429691-C3).

The TCP population in our previous dataset was most abundant at PCW12 and remained detectable at later stages, although its proportion relative to the total cerebellar cells gradually declined after PCW14. The decreased proportion of TCP cells among total cerebellar cells at later fetal stages may reflect the progressive transformation and absorption of the rhombic lip, along with the rapid expansion of other cerebellar structures. This conforms to the expected pattern for a transitional intermediate, whereas an artifact from contamination would be expected to exhibit a more random and uniform distribution over time. Debris would tend to reflect the most abundant mRNAs in the tissue, and there was no enrichment of housekeeping genes. Thus, the *in-silico* re-analysis of our single-cell transcriptomics of cerebellar samples, using more stringent QC measures and additional statistical tests, provide robust evidence re-confirming the existence of the TCP population in the human fetal cerebella. *It is important to note that the assertion that results obtained using commonly applied filtering criteria and quality control measures are merely artifacts fundamentally challenges the validity of any analysis employing similar methodologies*.

### Multiple orthogonal experimental validations for the existence of TCP cells by RNAscope *in situ* hybridization, Xenium spatial transcriptomics, and immunohistochemistry

#### Validation of TCP existence by RNAscope fluorescent multiplex in situ assays

In addition to the immunohistology validation of the presence of TCP cells in the cerebellar and rhombic lip ventricular zone at early developmental stages in our previous study^6^, we have further performed fluorescent mRNA *in situ* hybridization of TCP signature markers (*SOX11, HNRNPH1)*, along with *Protogenin* (*PRTG*), a recently identified rhombic lip stem-like cell marker^13^, and a UBC progenitor cell marker *EOMES*, on the midsagittal sections of human fetal cerebella at PCW11, PCW14 and PCW17 using RNAscope, a widely recognized gold standard for validating gene expression and spatially mapping^20^. Cells labeled with *SOX11* and *HNRNPH1*, were notably enriched in the rhombic lip, especially the rhombic lip ventricular zone (RL^VZ^) at the early developmental stage (**Fig. 2a-c**). At PCW11, cells expressing TCP signature genes are predominantly co-localized with *PRTG*^+^ stem-like cells in the RL^VZ^, with minimal overlap observed with *EOMES*^+^ UBCs at this early stage (**Fig. 2a**). At the later developmental stages PCW14 and PCW17, TCP cells appear to transition from co-localizing with *PRTG*^+^ cells to displaying increased spatial overlap with *EOMES*^+^ UBC cells in the RL^SVZ^ (**Fig. 2b, c**). These characteristics may represent the developmental state of TCP as transitional cells, that is, the transition from neural stem-like cells to differentiating neuronal progenitor cells. This expression pattern mirrors that of *PRTG* during early developmental stages near or prior to PCW11, suggesting that TCP progenitor cells may represent a downstream transitional population derived from *PRTG*^+^ stem-like cells and potentially serve as a candidate cell of origin for Group 3 (G3) medulloblastoma (MB)^13^. Moreover, the positional shift of TCP cells toward the RL^SVZ^, a primary source of UBCs, at later fetal stages aligns with our previous identification of TCP-like tumor cells in UBC-derived Group 4 MB^6^, suggesting that the transitional identity of TCP cells may represent a shared developmental intermediate across distinct MB subgroups. Notably, the combined expression of TCP signature markers enables precise identification of this transitional cell population. The use of multiple markers to delineate a distinct cell type or state is a widely accepted approach, ensuring specificity within complex and dynamic developmental contexts ^21,22^. Importantly, the strong and regionally enriched expression of TCP signature genes in the rhombic lip supports the selective localization and existence of the TCP cell population.

The *Matters Arising* stated that TCP marker genes display “*ubiquitous expression across all cell types*”, however, this is contradictory to their identification of the distinct TCP cell population after the reanalysis of our single-cell RNA-seq datasets (MA-Fig. 1). Importantly, the *Matters Arising* claimed “*RNAScope in situ hybridization analysis of proposed TCP marker genes (SOX11, HNRNPH1) across multiple stages of human cerebellar development (spanning PCW11-17) revealed diffuse expression of both genes in the cerebellum, with no regional enrichment*”. However, despite apparent chromogenic overdevelopment in the staining, the intense dark brown signal of TCP signature genes in the rhombic lip—distinct from the lighter red staining in adjacent germinal regions—clearly indicates enriched expression in the cerebellar rhombic lip at PCW11 (MA-Fig. 2f). More strikingly, TCP signature gene expression is robustly enriched in the rhombic lip at PCW14 and PCW17. These data are consistent with our RNAscope fluorescent *in situ* hybridization results (**Fig. 2**). This spatially restricted pattern across developmental stages contradicts the claim of nonspecific or ubiquitous expression and further supports our conclusions. Thus, contrary to their assertion, the selective and robust enrichment of TCP signature genes in the rhombic lip provides strong evidence for the existence of TCP cells, rather than refuting it.

#### Validation of the existence of TCP cells by Xenium spatial transcriptomic profiling in the human fetal rhombic lip

To independently validate the presence of TCP cell population, we performed high-resolution spatial transcriptomic analysis using the Xenium 5K platform on midsagittal sections of human fetal cerebellum at PCW 12.5, 14, and 16. Xenium is a next-generation *in situ* transcriptomics technology developed by 10x Genomics that enables multiplexed detection of thousands of genes with subcellular resolution while preserving the tissue’s spatial architecture. Compared to RNAscope *in situ* hybridization, which is based on a branched DNA amplification system and double-Z probe design for single-gene detection with high sensitivity, Xenium relies on barcoded probe pairs and provides broader transcriptomic coverage to generate high-throughput gene expression maps in a spatial context.

Xenium spatial transcriptomic profiling revealed distinct spatial localization of TCP signature genes *SOX11* and *HNRNPH1* within the rhombic lip, supporting the regional enrichment of this transitional progenitor population (**Fig. 3a–c**). Co-localized expression of the UBC neuronal progenitor marker *EOMES* and the rhombic lip progenitor marker *OTX2* within the *SOX11*^*+*^*/HNRNPH1*^+^ cell population further delineated the TCP cells in the rhombic lip progenitor zone, providing additional transcriptional context for the TCP niche in the rhombic lip. These Xenium spatial mapping data corroborate the RNAscope *in situ* hybridization findings (**Fig. 3a–c**), demonstrating consistent enrichment of TCP cells in the rhombic lip and increased spatial overlap with the rhombic lip subventricular zone (RL^SVZ^) marked by *EOMES*^+^ at later developmental stages (e.g., PCW14 and PCW16). Thus, these data further confirm the spatially and temporally restricted presence of TCPs in the human fetal rhombic lip and reinforce their identity as a transcriptionally distinct, intermediate population within the uniquely human developing cerebellar germinal zone.

**Figure 3.**
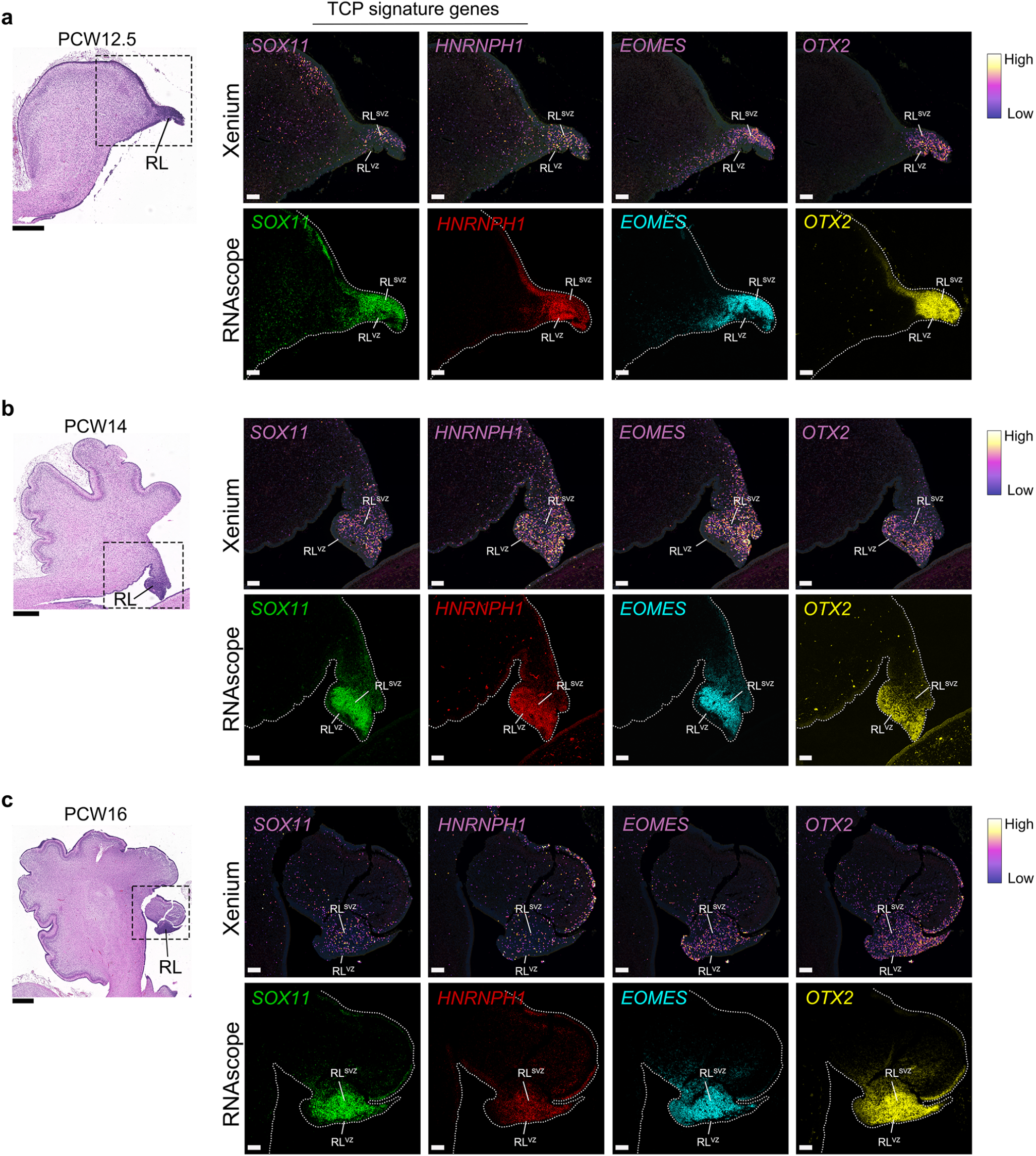
Xenium 5K spatial transcriptomics validates the existence of the TCP population in the rhombic lip in the human fetal cerebellum. **a**-**c**, Left, hematoxylin/eosin (H&E) staining of midsagittal sections in the human fetal cerebellum at post-conception weeks PCW12.5 (a), PCW14 (b), and PCW16 (c). Boxed areas: the rhombic lip (RL). Scale bars, 500 μm. Spatial gene expression maps by Xenium *in situ* (right, upper) and RNAscope fluorescent mRNA *in situ* labeling (right, lower panels) of TCP signature genes, *SOX11, HNRNPH1*, along with a UBC neuronal progenitor marker *EOMES* and rhombic lip progenitor marker *OTX2* in the rhombic lip at the developmental stages as indicated. Scale bars, 100 μm. RNAScope probes: *SOX11* (Advanced Cell Diagnostics, Cat#443871-C2), *HNRNPH1* (Advanced Cell Diagnostics, Cat#1283271-C1), *EOMES* (Advanced Cell Diagnostics, Cat# 429691-C3), and *OTX2* (Advanced Cell Diagnostics, Cat# 484581-C3).

#### Validation by immunohistochemistry in additional fetal cerebellar specimens

In our original study^6^, we validated the presence of TCP cells marked by SOX11 and HNRNPH1 co-expression in the cerebellar and rhombic lip ventricular zone using immunostaining at PCW12. By analyzing additional human fetal cerebellar tissues, we re-demonstrate the presence of the TCP population marked by SOX11 and HNRNPH1 in the rhombic lip of the cerebellar samples at PCW 12 (**Fig. 4a**). The TCP population, marked by high SOX11 and HNRNPH1 expression, is enriched in the rhombic lip ventricular zone and overlaps with a subset of SOX2^+^ stem-like cells, while remaining distinct from EOMES^+^ UBCs at this stage (**Fig. 4b**), consistent with our original study.

**Figure 4.**
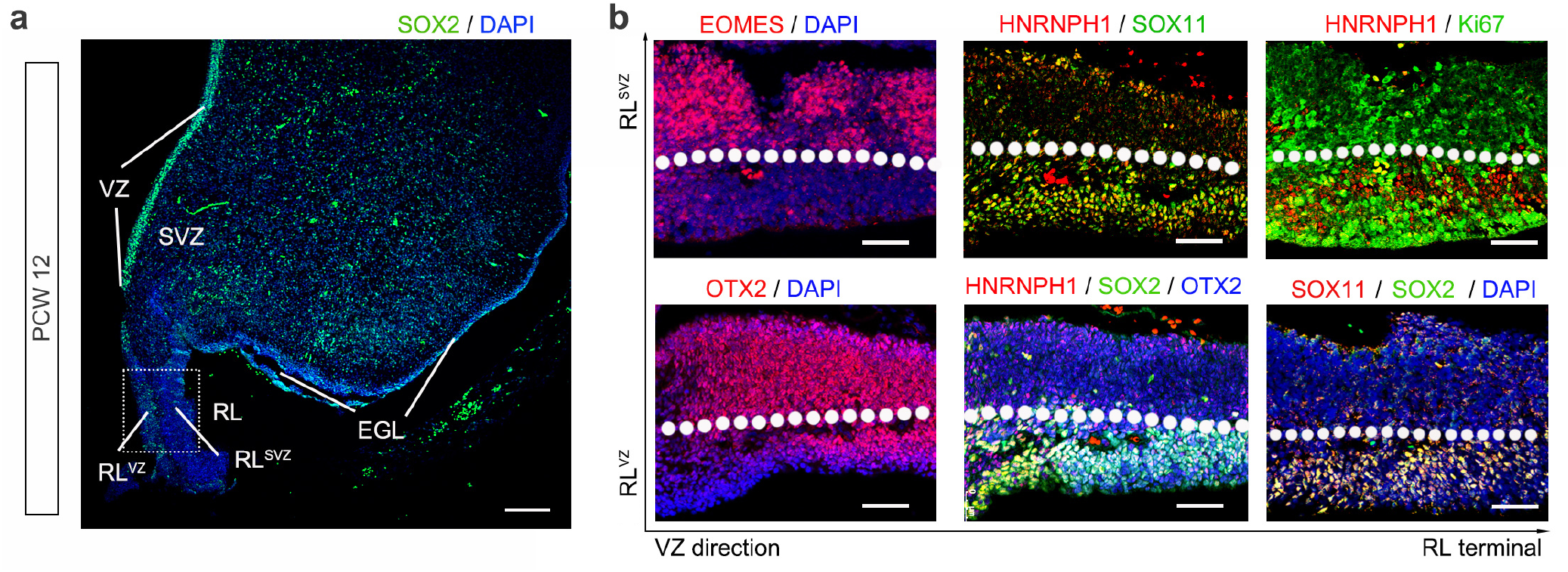
Immunohistochemistry of additional human fetal cerebella re-confirms the existence of the TCP population enriched in the rhombic lip. **a, b**, Representative images of the rhombic lip (RL) region of human fetal cerebella at PCW 12 immunostained for the markers as indicated. The boxed area in panel a is shown in high magnification in panel b. The dash lines in panel b show the boundary between RL^VZ^ (ventricular zone) and RL^SVZ^ (subventricular zone). Ki67, cell proliferation marker; SOX2, a stem-cell marker; OTX2, rhombic lip marker; EOMES, UBC marker. Scale bars, 500 μm in a, 100 μm in b.

Overall, multiple orthogonal lines of evidence, including RNAscope fluorescent multiplex in situ hybridization, Xenium spatial transcriptomics, and immunohistochemistry on additional specimens across developmental stages, reconfirm the presence of *SOX11*^*+*^*/HNRNPH1*^*+*^ TCP cells enriched in the rhombic lip, as reported in our original study. These findings support that this population represents a bona fide cell type rather than background noise. Together, these validation approaches using independent samples from distinct stages reaffirm the robustness of our original findings on the presence of TCP cells in specific germinal zones such as the rhombic lip during human fetal cerebellar development.

### TCP-like population identified across two independent human cerebellar datasets

The *Matters Arising* stated that transitional cerebellar progenitor (TCP) cells are absent in the recently published cerebellar single-cell dataset by Sepp et al. (2024)^9^ based on PCA projection analysis. It is worth noting that PCA projection of cell types from the Sepp dataset (derived from frozen nuclei) onto our dataset (derived from fresh whole cells) provides only predictive approximations rather than definitive conclusions. However, our independent analysis of exonic gene expression across cerebellar cell types within the Sepp dataset notably corroborates the presence of a TCP-like population. We analyzed the cerebellar single-nucleus transcriptomics dataset across cell types by Sepp et al.^9^, spanning PCW 7 to PCW 20 into adulthood. Since our whole-cell transcriptomic single-cell data primarily map to exonic regions derived from mature spliced mRNAs, we plotted the pseudo-bulk expression of exonic RNA transcripts based on the published website (https://apps.kaessmannlab.org/sc-cerebellum-transcriptome/). Exonic RNAs represent coding transcripts and closely correlate with protein expression, aligning well with our RNAscope mRNA *in situ* hybridization and immunohistology data. From the published Sepp dataset website, we observe that the exonic RNA expression of TCP signature genes such as *SOX11* and *HNRNPH1* is highly enriched in the cell type classified as “VZ_neuroblast” (ventricular zone neuroblasts) at PCW11 (**Fig. 5a**), a developmental stage closely aligning with PCW12, where we identified an enrichment of TCP cells. Although the nomenclature used for “VZ-neuroblast” cell type defined in the Sepp dataset differs from that in our previous classification, their strong expression of TCP signature genes suggests that these cells are analogous to the TCP population identified in our earlier study.

**Figure 5.**
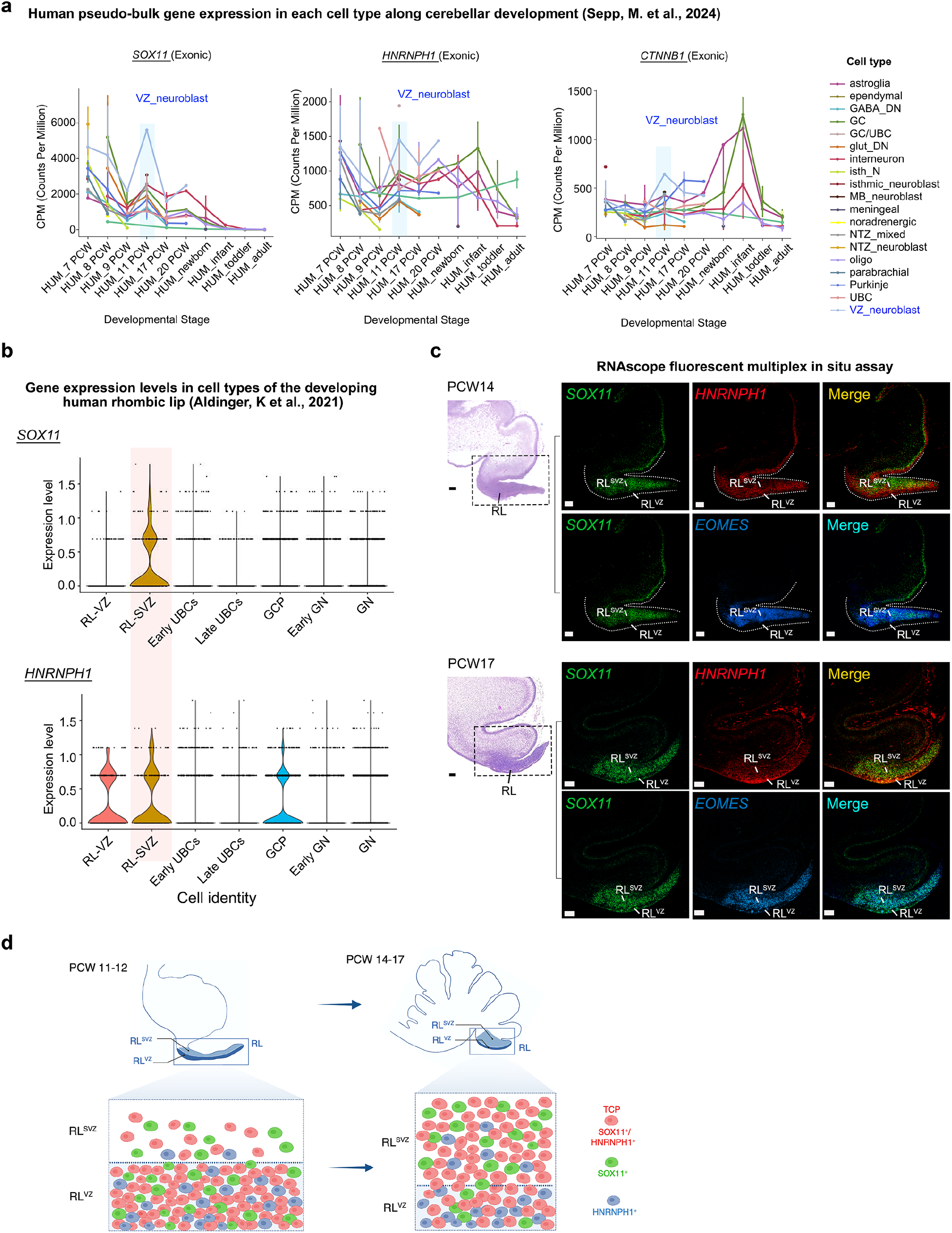
TCP-like cells identified across two independent human cerebellar single-nuclei transcriptomic datasets. **a**. Pseudo-bulk expression of TCP signature genes *SOX11, HNRNPH1*, and *CTNNB1* across different cell types and developmental stages defined in the Sepp study (2024)^9^. Notably, the high expression levels of these genes in VZ_neuroblast cells at PCW 11 are indicated. *SOX11* (exonic, ENSG00000176887), *HNRNPH1* (exonic, ENSG00000169045), *CTNNB1* (exonic, ENSG00000168036). **b**. Violin plots depicting the expression levels of TCP signature genes, *SOX11* and *HNRNPH1*, enriched in the rhombic lip (RL) progenitor cells such as RL-SVZ progenitors across the cell types of the developing human rhombic lip from different stages PCW 9, 10, 11, 14, 17, and 21 from the Aldinger dataset (2021)^8^. **c**, RNAscope fluorescent multiplex mRNA labeling of TCP signature genes (*SOX11, HNRNPH1*) with an SVZ-enriched UBC progenitor marker *EOMES* in the rhombic lip at PCW14 and PCW17 as indicated. Scale bars, 100 μm. **d**. Schematic diagram depicts *SOX11/HNRNPH1*-expressing TCP cells transition from the *PRTG*^+^ stem-like progenitor zone in the RL^VZ^ at early stages (e.g., PCW11) to the *EOMES*^+^ UBC-marked RL^SVZ^ at later stages (e.g., PCW14–17) during human fetal cerebellar development.

Consistently, analysis of gene expression levels in different cell types classified by another independent single-nucleus RNA-seq dataset from the developing human fetal cerebellum at multiple stages, previously published by Aldinger et al. (2021)^8^, also identified a population of TCP-like cells highly enriched for *SOX11* and *HNRNPH1* within the rhombic lip region such as the subventricular zone RL^SVZ^ (**Fig. 5b**). These findings are consistent with our RNAscope fluorescent multiplex *in situ* hybridization results, which demonstrate the presence of TCP cells marked by *SOX11* and *HNRNPH1* in the rhombic lip such as the RL^SVZ^ in the human fetal cerebellum (**Fig. 5c, d**).

Collectively, analyses of two independent cerebellar single-nucleus transcriptomic datasets reveal a ‘VZ-neuroblast’ population in the fetal cerebellum and a progenitor population in the rhombic lip subventricular zone, both of which transcriptionally resemble TCP cells, providing further support for the existence of the TCP population described in our original study.

### Deconvolution re-analyses confirm TCP-like population in malignant MB tumors

In our original study, we used CIBERSORTx deconvolution analysis^23^ using a set of the marker genes of each fetal cerebellar cell type, and showed the enrichment of the TCP signature in malignant MB tumors, especially the G3-MYC subgroup of MB^6^. We have further performed additional deconvolution analyses including MuSiC (MUlti-Subject SIngle Cell deconvolution)^24^ on the bulk MB tumor dataset using default parameters and our annotated scRNA-seq fetal cerebellar cell-type object. Even under these conditions, we detected the enrichment in a TCP-like population in malignant MBs. This population was especially prominent in G3-MYC^high^ tumors (**Fig. 6a**). Consistent with previous literature^14^, we also detected enrichment of GCP and UBC signatures specifically in SHH MB and G4 MB, respectively (**Fig. 6a**), validating the approach. Similarly, our analysis using the CIBERSORTx deconvolution approach^25^, with all the gene sets of expressed genes in each cluster, also revealed enrichment in the TCP-like signature in malignant MB tumors including G3-MYC^high^ tumors (**Fig. 6b**). Therefore, both MuSiC and CIBERSORTx deconvolution analyses using the gene set from each cell-type cluster yielded consistent results that showed enrichment of the TCP signature in malignant MBs. It is important to note that: 1) The analyses performed in the *Matters Arising* using different methods yielded inconsistent results (MA-Fig. 2g), and failed to reproduce previously established similarities between GCPs and SHH MB, or between UBCs and G4 MB^14^, which serve as positive controls. These discrepancies raise concerns about the validity of their computational approaches. 2) Methods of deconvolution of bulk RNA-seq data are only predictive of potential cell populations in tumor tissues, whereas our scRNA-seq transcriptomics analysis at single-cell resolution clearly identified the TCP-like population in individual G3 tumors (**Fig. 6c, d**). Thus, rather than relying on deconvolution algorithms to infer tumor cell origins, our unbiased single-cell transcriptomic analysis of MBs—supported by immunohistochemistry and single-cell proximity ligation assays—identified a transcriptionally similar TCP population in the tumor tissues^6^, establishing a biologically relevant link to their developmental counterparts.

**Figure 6.**
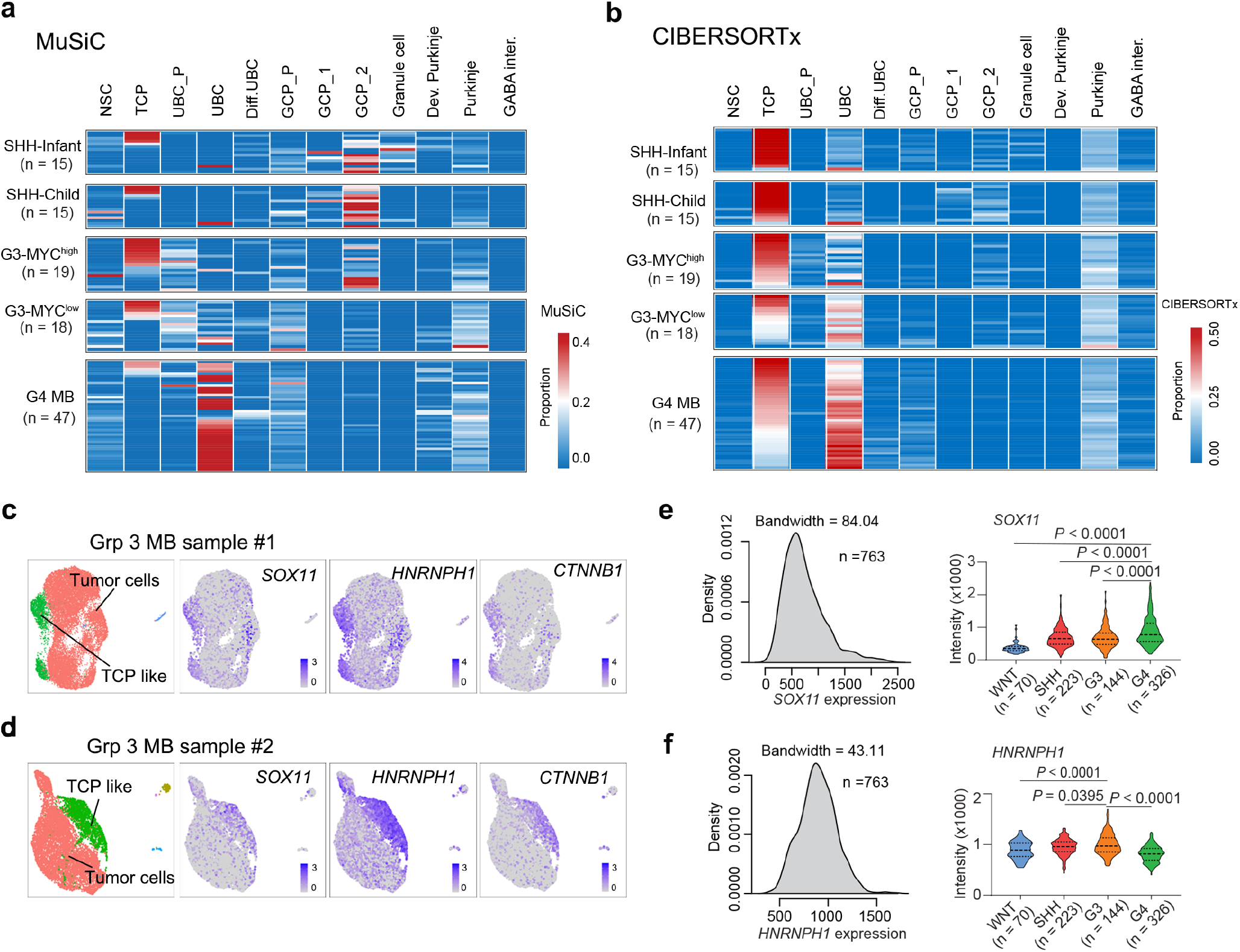
Additional deconvolution analyses validate the presence of TCP-like tumor cells within malignant MBs. **a**-**b**, Heatmap showing the predicted cell-type proportions in individual MB patients using a) MuSiC or b) CIBERSORTx deconvolution. Single-cell expression matrices of all gene sets generated by the Seurat program in each cluster were used for MuSiC or CIBERSORTx deconvolution analyses. For the bulk RNA-seq matrix in MB tissues, we used 13,889 genes (FPKM >1) from the entire gene list. **c**-**d**, Representative UMAP of cell-type compositions (left) and expression distribution of TCP marker genes in the UMAP plot (right) for two independent G3 MB tumor samples. **e**-**f**, Density plots of expression of e) *HNRNPH1* and f) *SOX11* in the normalized bulk microarray dataset of 763 MB tissues^26^ (left) and the violin plots of expression of these genes in tumor groups. Significance was determined by one-way ANOVA followed by Tukey’s multiple comparison tests.

In our original study, we compared the expression of TCP signature genes *HNRNPH1* and *SOX11* in MB subgroups using a normalized bulk microarray expression dataset from MBs (n = 763)^26^, which approximated a normal distribution^27^ (**Fig. 6e, f**). We therefore performed statistical analyses using the actual expression values^26^. An unpaired *t*-test showed that TCP signature genes *HNRNPH1* and *SOX11* were expressed at significantly higher levels in G3 and G4 MBs, respectively, than in other subgroups in our original study. By using additional ANOVA analysis with Tukey’s multiple comparisons, a rigorous statistical method for multiple group comparison, we reached the same conclusions (**Fig. 6e, f**). In contrast in the *Matters Arising*, Limma was used for multiple testing correction (MA-Fig. 2h), which is inappropriate in this context as it overestimates the number of tests performed. This is particularly problematic given the limited number of genes and cohorts analyzed. Therefore, adjusting the p-values based on the total number of genes is unwarranted.

The intellectual disability gene *SOX11* is expressed in the cerebellum and medulloblastoma, and its mutations have been implicated in human neural developmental disorders^28,29^. Similarly, mutations in *HNRNPH1* are linked to a distinct syndromic neurodevelopmental syndrome^30^, characterized by recurrent cerebellar structural abnormalities. *HNRNPH1* functions as an essential gene in 88% of tested cell lines according to DEPMAP^31^, consistent with our original finding that it is critical for the growth of aggressive MBs, such as Group 3 MB.

## Conclusion

Our re-analyses using more stringent filtering criteria and quality control measures, along with multiple orthogonal experimental validations—including RNAscope in situ hybridization, Xenium-based spatial transcriptomics, and immunohistochemistry in additional fetal cerebellar samples across developmental stages—reconfirm the presence of the TCP population in the human fetal cerebellum, particularly in regions such as the rhombic lip, as reported in our original study^6^. In addition, the independent identification of the “VZ-neuroblast” and “RL-SVZ” populations resembling TCP cells across two cerebellar single-nucleus transcriptomic datasets^8,9^ further supports the existence of the TCP population described in our original study. Our additional quality control evidence provided here shows that TCP cells were not detected due to “liberal” filtering criteria, ambient RNA contamination or empty/damaged droplets. Importantly, we provide RNAscope fluorescent multiplex *in situ* hybridization and high-resolution Xenium spatial transcriptomics for the existence of the TCP cells in specific geminal zones of the human fetal cerebellum such as the rhombic lip across different fetal developmental stages that we proposed in our original study. Thus, our findings are not an anomalous unique situation seen in a single donor. *It is important to note that differences in the use of fresh versus frozen cerebellar tissues, along with variability in the anatomical regions sampled, can impact cell-type clustering and resolution, and gene expression profiles, potentially contributing to discrepancies observed across single-cell transcriptomic studies*^*32-37*^.

Our multiple orthogonal validation approaches using RNAscope *in situ* labeling, Xenium spatial transcriptomics, and immunohistochemistry across multiple donors confirm the anatomical specificity of the TCP population—such as its localization to the rhombic lip—and support that it is not a bioinformatic artifact. Furthermore, we detect the TCP-like population in multiple malignant MB tumor tissues^6^. Thus, the consistent identification of TCP populations in both developing fetal cerebellar tissues and malignant MB samples strongly supports the biological validity of the originally described TCP population.

TCP cells in the rhombic lip may represent a transitional population that progresses and differentiates along the rhombic lip lineage trajectory, bridging early stem-like progenitors and committed neuronal precursors (**Fig. 4d**). At early fetal stages (e.g., PCW11–12), *SOX11*/*HNRNPH1-* expressing TCP cells are predominantly localized within the ventricular zone of fetal cerebella and the *PRTG*-expressing stem-like progenitor zone in the RL^VZ^. By later fetal stages (e.g., PCW14 and PCW17), a population of these cells appears to transition from the RL^VZ^ to the RL^SVZ^, a region marked by *EOMES*^+^ UBC cells (**Fig. 4d**). This stage-specific spatial distribution of TCP cells across distinct progenitor domains likely reflects their dynamic progression within the rhombic lip, transitioning from neural stem-like progenitors in the RL^VZ^ to differentiating neuronal lineages in the RL^SVZ^ during human fetal cerebellar development, in line with the model proposed in our original study. While *SOX11* and *HNRNPH1* are detectable in other cerebellar cell types, our analyses show that these TCP signature genes are highly enriched in the cerebellar ventricular zone and rhombic lip region during critical developmental windows of the human fetal cerebellum. Thus, reanalysis of our single-cell transcriptomic data with more stringent quality control, combined with *in-silico* interrogation of independent fetal cerebellar single-cell datasets and additional orthogonal validations of gene expression by RNAscope and Xenium spatial transcriptomics across multiple developmental stages, collectively provides robust and convergent evidence supporting the existence of the TCP population in the developing human fetal cerebellum.

## Acknowledgements

The authors would like to thank Drs. Charles Stiles, Joseph Powell, William Weiss, Richard Gilbertson, Martine Roussel, Huda Zoghbi, William Dobyns, Alexandra Joyner, Mario Suva, Jiyang Yu, Saulnier Olivier, Tamra Ogilvie, Kenny Campell, and Yi Zheng for discussions and comments. The illustrative diagram was created using BioRender.com.

## Methods

### Human fetal cerebellar and tumor tissues

Human fetal and tumor tissues were obtained from the Children’s Hospital of Fudan University and the Obstetrics and Gynecology Hospital of Fudan University. Written informed consent for the use of tissues in research was obtained from donors or the patients’ parents. The collection of fetal tissues was approved by the Institutional Review Board of the Obstetrics and Gynecology Hospital of Fudan University and Guangzhou Women and Children’s Medical Center.

### RNAscope *in situ* hybridization and immunohistochemistry

RNAscope fluorescent in situ hybridization assays were performed using commercially available probes from Advanced Cell Diagnostics, following the manufacturer’s instructions. The in situ hybridization probes used were: *SOX11* (Cat# 443871-C2), *HNRNPH1* (Cat# 1283271-C1), *PRTG* (Cat# 1096421-C3), *EOMES* (Cat# 429691-C3), and *OTX2* (Advanced Cell Diagnostics, Cat# 484581-C3). The immunostaining procedures followed the method as previously described^6^. For immunohistochemistry, we used the following primary antibodies: mouse anti-SOX2 (Santa Cruz Biotechnology, Cat# sc-365964), rabbit anti-SOX11 (Sigma, Cat# HPA000536; Thermo Fisher, Cat# 14-9773-82), rabbit anti-HNRNPH1 (Abcam, Cat# ab154894; Bethyl Laboratories, Cat# A300-511A), rat anti-EOMES (Invitrogen, Cat# 14-4875-52), and anti-OTX2 (Proteintech, Cat# 13497-1-AP).

### Xenium-based spatial transcriptomics

Freshly collected cerebellar tissue was immediately immersed in 10% neutral buffered formalin (NBF), fixed at 4°C for 16–48 hours, and subsequently embedded in paraffin. Formalin-fixed paraffin-embedded (FFPE) sample preparation and sectioning (5 μm thickness) were performed according to the 10x Genomics User Guide (CG000578). Deparaffinization and decrosslinking were conducted following the 10x Genomics protocol (CG000580). Xenium Prime in situ analysis was performed using the Xenium Prime 5K Human Pan-Tissue and Pathways Assay Kit (10x Genomics, PN-1000671), following the experimental guidelines in the 10x Genomics User Guide (CG000760). Briefly, tissue sections were hybridized with target-specific probes at 50°C for 16–24 hours. Subsequent washing, probe ligation, and amplification steps were carried out using the Xenium Slides & Sample Prep Reagents Kit (10x Genomics, PN-1000460). Amplified transcripts were stained using the Xenium Cell Segmentation Staining Reagents (10x Genomics, PN-1000661) for cell segmentation. Prior to imaging, sections were treated with autofluorescence quenching reagents and stained with DAPI. Fluorescence detection and transcript decoding were performed on a 10x Xenium Analyzer (10x Genomics, PN-1000481). Following Xenium detection, hematoxylin and eosin (H&E) staining was carried out according to the Post-Xenium Analyzer H&E Staining User Guide (CG000613).

### Human fetal cerebellar single-cell RNA-seq data processing

FASTQ files^6^ were aligned using CellRanger v3.1. Additional quality control metrics for each sample were computed using the established pipelines in CellRanger and Seurat as well as such as DoubletFinder^10^, SoupX^11^, and DropletQC^12^ with default or specified parameters.

### Deconvolution of MB tumors

Estimation of cell-type proportions within public CBTTC cohort expression profiles (https://cbtn.org/) was conducted using a website-based deconvolution approach CIBERSORTx (https://cibersortx.stanford.edu/) and MuSiC (https://github.com/xuranw/MuSiC) run on default parameters. MB bulk RNA-seq expression value (PPKM) and average expression (signature gene matrix) of each cell type from human fetal cerebellum were used as inputs in CIBERSORTx. MB bulk RNA-seq FPKM expression values and Seurat objects of neural stem cells and neuronal lineage cell types from the human fetal cerebellum were used as inputs for MuSiC.

## Code availability

Custom code is available at https://github.com/Luostlac/public1 and https://github.com/jumphone/public.

